# S100A4 inhibits cell proliferation by interfering with the RAGE V domain-S100A1

**DOI:** 10.1101/391136

**Authors:** M I Khan, T Yuan, R H Chou, C Yu

## Abstract

The Ca^2+^-dependent human S100A4 (Mts1) protein is part of the S100 family, and the S100A1 protein is the target of S100A4. Here, we studied the interactions of S100A1 with S100A4 using nuclear magnetic resonance (NMR; 700 MHz) spectroscopy. We used HADDOCK software to model S100A4 and S100A1, and we observed that S100A1 and the RAGE V domain have an analogous binding area in S100A4. We discovered that S100A4 acts as an antagonist among the RAGE V domain and S100A1, which inhibits tumorigenesis and cell proliferation. We used a WST-1 assay to examine the bioactivity of S100A1 and S100A4. This study could possibly be beneficial for evaluating new proteins for the treatment of cancer.

## 1. Introduction

The family of human S100 proteins are Ca^2+^-dependent, slightly acidic proteins comprising more than 20 family members with molecular weights of 9-13 kDa in vertebrates (1). There is increasing evidence of the scientific importance of many parts of S100 proteins. The disorder of these proteins, which contain malignant neoplasms, has been found repeatedly in many human diseases. A few of them have been proposed as medicinal aims or predictions of a therapeutic answer (2–4). Their observational interactive roles include the regulation of protein phosphorus, enzyme action, cell development and discrimination; many S100 proteins exhibit chemotactic and neurotrophic activities (2, 5, 6). Several S100 proteins are known to be possible markers of various cancers such as breast and colorectal cancer pancreatic, thyroid, gastric bladder, and melanoma (2). These proteins are generally found as a bunch on the human chromosome 1q21 (7). The family of EF-hand Ca^2+^-holding proteins is familiar to science, but intracellular Ca^2+^ mediates signals in an unknown fashion (8, 9). The S100 family has an insecure hydrophobic pocket that facilitates interactions of the protein (10–12). These proteins have the tendency to form hetero-dimers and homo-dimers (13) and appear to create many kinds of hetero-oligomers and homo-oligomers (14, 15). Oligomerization is a pathway of cell surface receptors for activation (16).

The S100A4 protein (also known as Mts1) is a part of the S100 superfamily, which includes the leading EF-hand Ca^2+^-holding proteins and regulates many proteins engaged in various cellular functions such as apoptosis, differentiation, proliferation, two-calcium ion (Ca^2+^) homeostasis, and energy metabolism (17–19). The S100 superfamily controls a large variety of essential cellular developments via protein-protein connections (8). Calcium binding EF-hand motif proteins have the tendency to undergo conformation changes of the destination protein by highlighting the hydrophobic areas in S100 proteins (20). EF-hand motif calcium binding initiates the action of the S100 proteins with structural changes and allows them to interact via selectivity (21, 22). The S100A4 protein was first deduced from stromas and tumors. In solution, the S100A4 protein takes the form of a homo-dimeric and acts as a metastasis-supporting protein (23, 24). The presence of S100A4 has now been demonstrated in many cancers (e.g., pancreatic gastric, colorectal, bladder, and breast). The S100A4 protein acts as a part of angiogenesis and tumor establishment (23–25). The EF-hand hinge area and the C terminus of the Mts1 protein are specifically related to another S100 protein. However, the majority of S100 proteins are related to target protein-protein binding. Calcium ion binding results in conformational changes in proteins to expose the hydrophobic pocket in helices 3 and 5 of the C-terminal EF-hand and the hinge region (26–28).

S100A1 is a part of the S100 family—it is expressed the most in cardiomyocytes (29). S100A1 has been noted in the heart, brain, skin, ovaries, thyroid gland, breasts, salivary glands, skeletal muscles, and kidneys. It is the source of various endometrial cancers such as melanoma, breast, thyroid, renal, endometrioid, and it is responsible for neurodegenerative disorders (29, 30, 39–43, 31–38). Due to the helix 3 and 4 conformation, S100A1 creates a large hydrophobic region between this helix, and many Ca^2+^-dependent target protein interactions take place in this region (44). Previously studies have demonstrated the interaction of S100A1 with other proteins such as ATP2A2, RyR1, TRPM3, RyR2, and RAGE (45–50). The conformational changes or activities of S100A1 support particular physiological roles. The S100A1 protein plays a crucial role in gene therapies, and it was recently used in human clinical trials related to heart failure (51).

We studied S100A4 as an inhibitor—it blocking the interface of the V domain and S100A1. All the details about S100A1-V domain interaction is reported by our group (52). This results, based on WST-1 assays, suggest that the inhibition properties arise from the bioactivity of S100A4 (53). We also present putative models of the S100A4-S100A1 complex. This study could possibly be beneficial for the development of new proteins useful for the treatment of cancer.

## 1. Materials and Methods

### 2.1 Materials

Ninety-nine percent ^15^NH_4_Cl (^15^N-ammonium chloride) and 99% D_2_O (isotopic-labeled deuterium oxide) were purchased from Cambridge Isotope Laboratories. All of the buffers and solutions were prepared using milli-Q water. The buffers for the NMR spectroscopy sample were filtered using a 0.22-μm antiseptic filter. The SW480 assay cells were bought from CCL-288 (American Type Culture Collection.

### 2.2 Expression procedures of S100A4 and S100A1

The S100A4 protein includes four cysteine residues (free). We used dithiothreitol (DTT) as a reductant in the buffer for the NMR experiments. The cDNA of the S100A4 was obtained from Mission Biotech Company. The S100A4 protein contains the vector pET21b, and *E. coli* was used for the transformed and over-expressed in the host cell BL21 (DE3). M9 medium was used for bacterial growth, and ^15^NH_4_Cl was the source of ^15^N labelled. The cultures were grown at a temperature 37°C until they obtained an optical density (OD) of 0.75–1.00. 1.0 mM of IPTG used to induce the culture, which was grown at 25°C for 15–18 h. The cells were lysed and harvested using a sonicator in buffer composition (300 mM KCl, 1 mM DTT, 2 mM CaCl_2_, 20 mM Tris, 1 M (NH_4_)_2_SO_4_ and 1 mM EDTA at pH 7.5). The sample was sonicated again for 30 min and centrifuged at 11000 rpm. S100A4 was obtained in soluble form, and purification was carried out using a HiPrep 16/60 Phenyl FF column with hydrophobic interaction chromatography (HIC, GE Healthcare. The HIC column was rinsed with buffer-1 containing 300 mM KCl, 1 mM DTT, 2 mM CaCl2, 20 mM Tris, 1 M (NH4)2SO4 and 1 mM EDTA at pH 7.5 and then washed with buffer-2 containing 1 mM EDTA, 20 mM Tris, 300 2 mM CaCl_2_, mM KCl, and 1 mM DTT at pH 7.5. Finally, it was eluted with a buffer containing 10 mM EDTA, 20 mM Tris, and 1 mM DTT at pH 7.5. The S100A1 portion was placed in the buffer with 20 mM Tris-HCl, 1 mM DTT and 0.02% NaN_3_ at pH 7.5 and was purified with a Q-Sepharose column with Hi-Prep Phenyl FF 16/10 on a AKTA FPLC system. S100A4 was eluted by a gradient with a buffer containing 1 mM DTT, 20 mM Tris-HCl, 0.02% NaN_3_ and 1.5M NaCl at pH 7.5. The S100A4 protein was exchanged with an NMR buffer containing 8 mM NaCl, 0.1 mM EDTA, 16 mM Tris, 6 mM CaCl_2_, and 0.34 mM NaN_3_ at pH 6.0 using a Millipore centrifuge tube. The purity of S100A4 was confirmed using SDS-PAGE (Fig S1).

The cDNA of S100A1 (1–93 amino acids) was inserted into XhoI or NdeI restriction sites. The expression vector pET20b was used for cloning, and host cell Rosetta™ (DE3) was used for the transformation and expression. Details of the purification of the S100A1 protein have been previously reported (54). The purity of S100A1 was identified using SDS-PAGE (Fig S2).

We first dissolved S100A1 and S100B in 8M urea separately to make them to be monomer. Then we mixed them together to generate mixture of S100A1 dimer (25%), S100B dimer (25%) and heterodimer S100A1-S100B (50%). However, it is difficult to separate these three species,

### 2.3 ^1^H-^15^N HSQC NMR titration

A Varian 700 MHz NMR cryogenic spectrometer was used for the HSQC titrations at 25°C. All of the samples for the titration contained the same buffer at pH 6.0: 0.1 mM EDTA, 8 mM NaCl, 6 mM CaCl_2_, 16 mM Tris, and 0.34 mM NaN_3_. The HSQC titrations were carried out by adding S100A1 to the ^15^N S100A4 in proportions of 1:0 and 1:1 molar. A second titration was also performed with ^15^N labeled S100A1 by adding S100A4 in proportions of 1:0 and 1:1 molar. The NMR HSQC spectra were purposely overlapped to determine whether the intensities should be decreased or the cross peaks shifted.

### 2.4 Docking study of S100A1 and S100A4 (HADDOCK)

The HADDOCK program (version 2.2) was used to create the structure of the S100A1-S100A4 complex. The structures of S100A1 and S100A4 were selected from the PDB (ID: 2MRD, 2LP3, separately). The docking study and successive refinement were carried out using a number of ambiguous-interaction restraints (AIRs) and residues that represented the HSQC keyed-out peaks with decreased intensities or considerable chemical shifts (52). The reported S100A1-S100A4 complex was selected from the cluster with the lowest energy, and the complex was illustrated and displayed using software called PyMOL (55).

### 2.5 Dissociation constant measurements from isothermal titration calorimetry (ITC)

Dissociation constant was measured by using Microcal iTC200 Calorimeter on the basis of heat change throughout the titration of S100A1 with S100A4. For ITC measurements, we used a buffer containing 0.1 mM EDTA, 8 mM NaCl, 0.34 mM NaN3, 6 mM CaCl2, and 16 mM Tris at pH 6.0. S100A1 and S100A4 centrifuged and degassed with the help of vacuum. Titration was carried out at 25^0^C by injecting 4 μL of S100A4 as ligand protein (25 times; 2mM concentration) into 0.2 mM of S100A1. Finally, buffer-protein control was used to correct the titration curve and the Origin software (provided by Microcal) was used to analyzed the data.

### 2.6 Dissociation constant measurements based on fluorescence

For binding constant measurement of protein-ligand and protein-protein interaction, fluorescence is a widely used method (56–58). A F-2500 fluoro-spectrophotometer (Hitachi) was used for the experiments. In S100A4, there is no tryptophan. The presence of the tryptophan residue in the S100A1 protein excites and causes the protein to fluoresce. For S100A1, the excitation frequency of the tryptophan absorption band was found at 295 nm, and wavelength of 295 nm was used for the excitation. At a wavelength of 351 nm, S100A1 exhibits an emission band. Over the range of 305–404 nm, we noted the emission wavelengths. For the fluorescence measurements, we used a buffer containing 0.1 mM EDTA, 8 mM NaCl, 0.34 mM NaN3, 6 mM CaCl2, and 16 mM Tris at pH 6.0. The S100A4 concentrations were increased from 0 to 24 μM in increments of roughly 0.80 μM to the S100A1 which consist of 3.50 μM concentration. Variation in emission spectra was observed as the S100A4 concentration was increased in the complex solution. The curve was fitted and the following equations were used to calculate the dissociation constant of the S100A1-S100A4 complex (59).

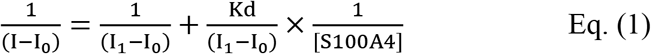

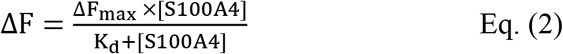

In Equation 1, I_0_ refers to the intensity of the free S100A4 in the solution; I and I_1_ refer to the emission intensities at the middle and maximum concentration of S100A4, respectively. F is the fluorescence intensity change between I_0_ and I. F_max_ is the maximum fluorescence difference and Kd is the dissociation constant. The actual curve was processed further as curve fitting and slope was calculated to get the Kd.

### 2.7 Bioactivity study with the WST-1 assay

We used the WST-1 assay to determine the physiological condition of S100A1 proteins. In living cells, WST-1 is split to a soluble formazan via mitochondrial dehydrogenases. The number of formazan created is directly proportional to the dehydrogenase enzymatic activity. Therefore, the difference in OD values at a suitable wavelength is correlated with the number of active metabolically cells in the culture. The WST-1 assay was conducted in the same manner as the Roche method. We placed the SW480 cells at a density of 5×10^3^ cells/well in a 96-well plate one day prior to the experiments. Next, in a serum medium consisting of bovine serum albumin (0.1%), the cells were incubated for one day. The serum-starved cells were titrated with 100 nM S100A1 with S100A4 and left free of S100A4 protein for an additional 48 h to detect the exchange of the proportional cell number. Prior to collecting the 1/10 volume of the WST-1 chemical agent was added to all of the wells. This agent consisted of 100 μL of culture, and the cells were incubated for an additional 4 h at 37°C. The culture cell plate was then incubated with slight agitation for 10 min on an incubator. The absorbance was determined at a wavelength of 450 nm using a synergy 2 micro-plate reader (52).

## 3. Results

### 3.1 The binding site of S100A4 and S100A1

^1^H-^15^N NMR HSQC is generally used to determine the binding site between a ligand and a protein. Interacting residues of the S100A4 and S100A1 proteins can be determined by calculating the resonances on the HSQC NMR spectra of S100A4 and correlated them with S100A4 in the complex with S100A1. Superimposed HSQC NMR spectra of free ^15^N S100A4 and the ^15^N S100A4-S100A1 complex are shown in Fig 1A. Portions of the NMR signals were decreased after the addition of S100A1 to ^15^N S100A4. The NMR HSQC signals of an S100A4 protein complex with S100A1 residues at the interaction site were lower than those acquired with free S100A4. The assignment of S100A4 (BMRB under accession number 25069) was dome by our group (60). This finding is likely due to the affected nuclei at the interface between the S100A4 and S100A1 proteins.

**Fig 1.**
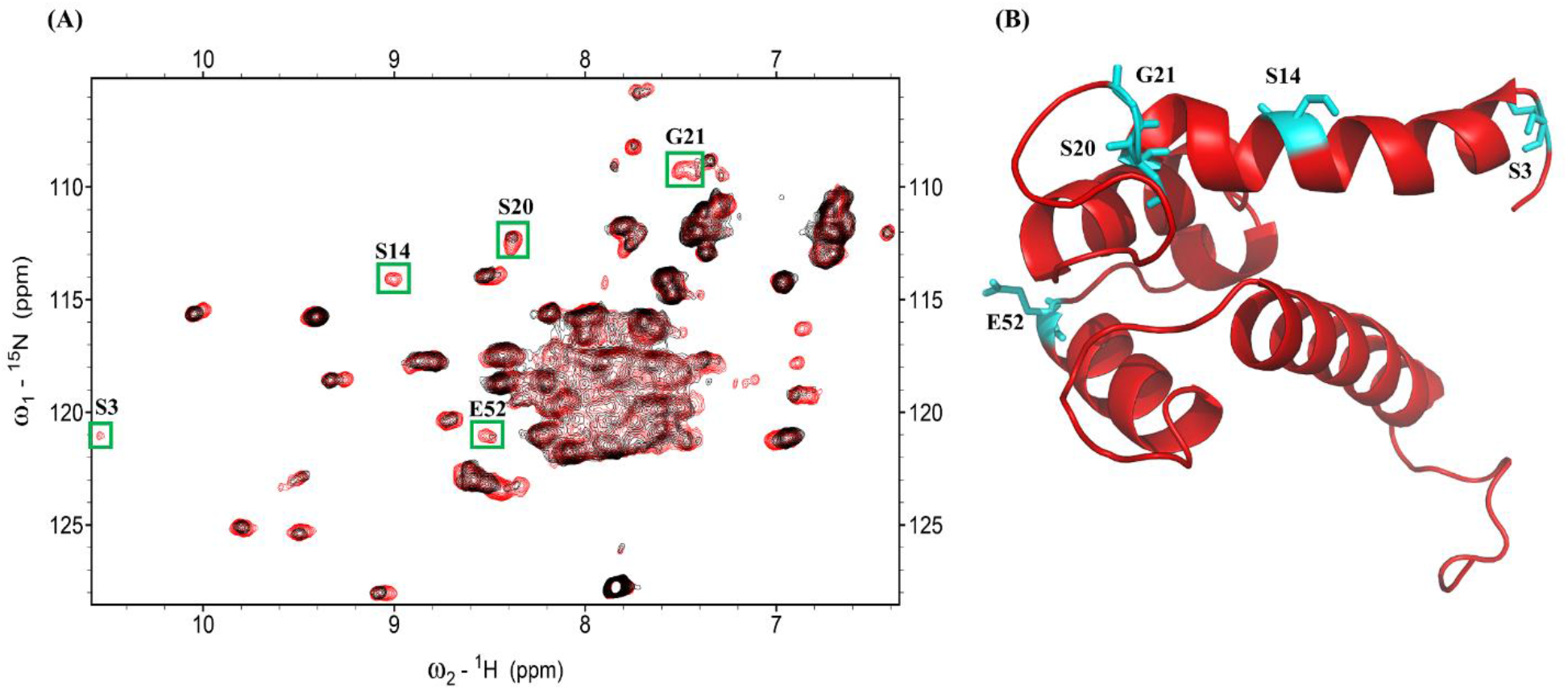
(A) Superimposed HSQC of the ^15^N S100A4-S100A1 complex (red) and ^15^N S100A4 (black). Peaks indicating reduced intensity appear in green boxes. (B) Ribbon diagram showing the structure of S100A4; decreasing residues are labeled on the structure in cyan.

The adjacent nuclei at the interacting site were reformed by another protein, which brought together and led to a decrease in the strength of peaks in the HSQC spectrum. When the S100 protein formed a complex with its target protein, the residues of the S100 protein were affected. Previous NMR HSQC studies noted changes in HSQC resonance among protein complex interfaces. Therefore, the spectra of free S100A4 and the S100A4-S100A1 complex were superimposed to classify the decreased intensity of S100A4 residues (S3, S14, S20, G21, E52) at the interface.

### 3.2 The interface site of S100A1 and S100A4

We also performed a reverse titration experiment on S100A4 titration with ^15^N S100A1 to define the residues of S100A1 binding to S100A4. The HSQC spectra of free S100A1 (The assignment of S100A1 in BMRB Entry 18101) (54) and S100A1 complexed with S100A4 (Fig 2A). Residues that exhibited a low intensity were selected for plotting on a cartoon model of S100A1 (Fig 2B): G1, E3, N13, H16, G43, F44, D52, A53, V76, L77, and A79.

**Fig 2.**
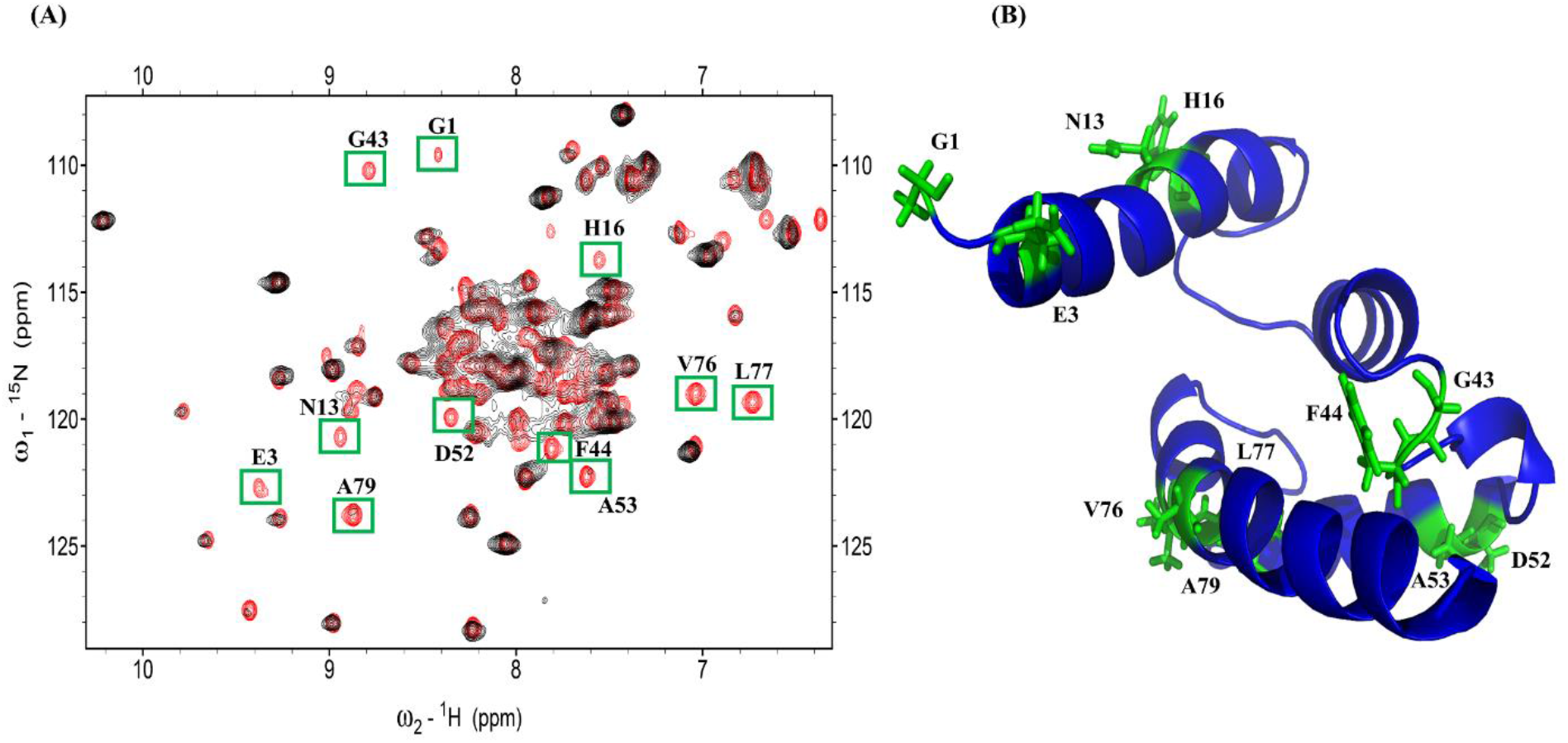
(A) Overlapping ^15^N S100A1 HSQC spectra (black) and the ^15^N S100A1-S100A4 complex (red). The cross peaks, which exhibited decreased intensity, are boxed in green. (B) Ribbon diagram showing the structure of S100A1—decreasing residues are labeled on the structure in green.

### 3.3 The S100A1-S100A4 complex structure

The complex structure of S100A4 and S100A1 was developed using the HADDOCK program to compute the protein complex. Ambiguous interaction restraints were developed based on the difference in resonance between the NMR HSQC of S100A4 and S100A1. Input parameters for the HADDOCK program were selected based on those residues that exhibited perturbations from the spectra (NMR HSQC). We also observed that some of the residues disappeared at the N-terminal of S100A1, which may be due to the flexibility and mobility of the N-terminal being high when a complex is formed.

Based on the HADDOCK results, we generated a heterodimeric complex of S100A4 and S100A1. The three-dimensional S100A1 and the S100A4 structures were obtained from PDB (ID: 2MRD,2LP3). Nearly 5000 complex structures were produced by applying rigid-body minimization with HADDOCK. The top 200 structures with lower energies after water refinement were used in this study. The structure of the heterodimer S100A1-S100A4 complex is shown in Fig 3.

**Fig 3.**
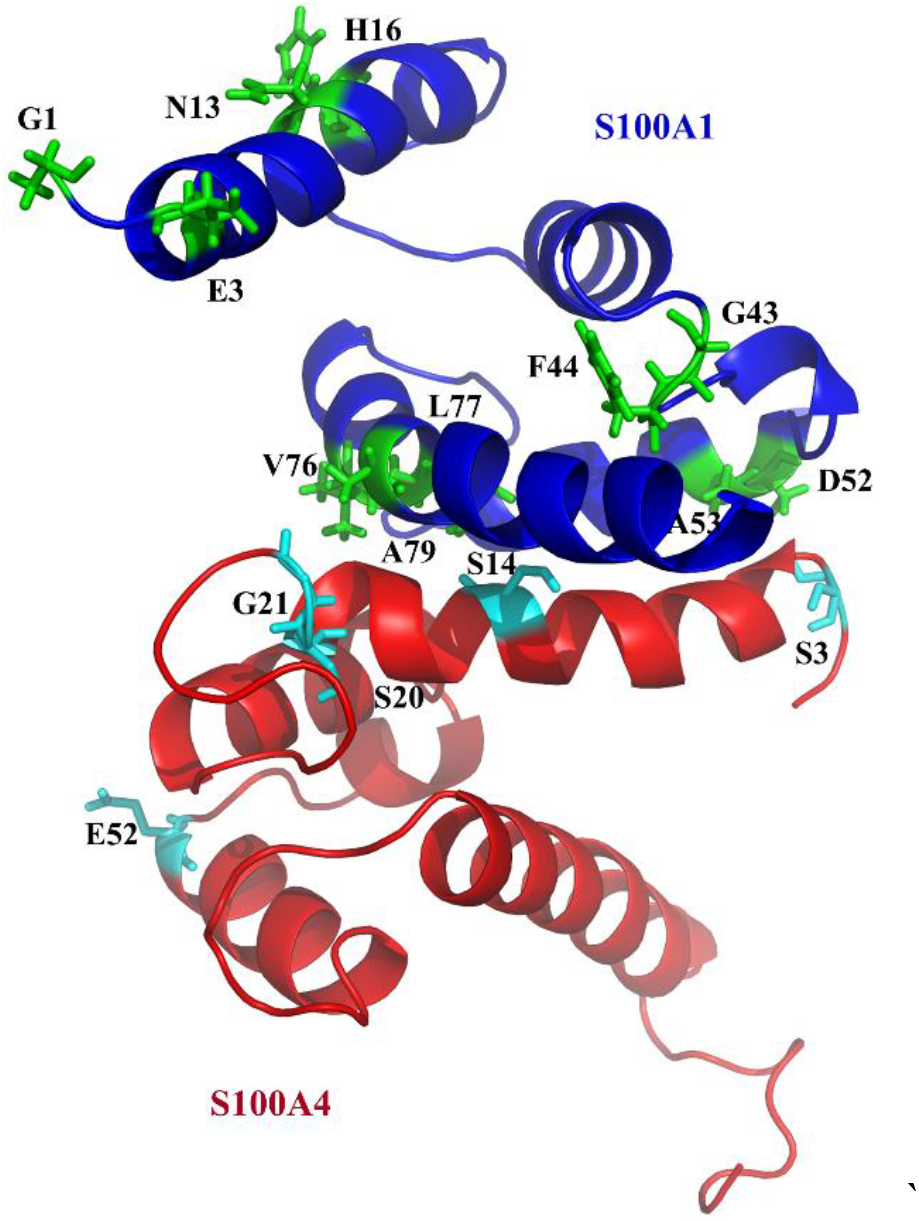
Model of the S100A1-S100A4 complex. S100A1 and S100A4 are shown in blue and red, respectively. The interacting residues are shown in cyan and green.

The Rampage plot analysis of the acquired complex S100A1-S100A4 had fair bond angles ψ and φ in the stereochemistry of the proteins. The Rampage plot (Fig S3) reveals that 91.6% of the complex residues were glycine free in the favorite region. Six residues (3.2%) were glycine free in the outlier area.

### 3.4 Dissociation constant measurements from isothermal titration calorimetry of the complex S100A1-S100A4

Isothermal titration calorimetry (ITC) is commonly used to study protein interactions. Isothermal titration calorimetry can measure the dissociation constant kd and changes in energy reliably; these changes in energy arise from protein-protein interactions (61). Isothermal titration calorimetry reveals evidence of binding sites of the protein and ligand. We observed a k_d_ 6.37 μM for the S100A1-S100A4 complex (Fig 4).

**Fig 4.**
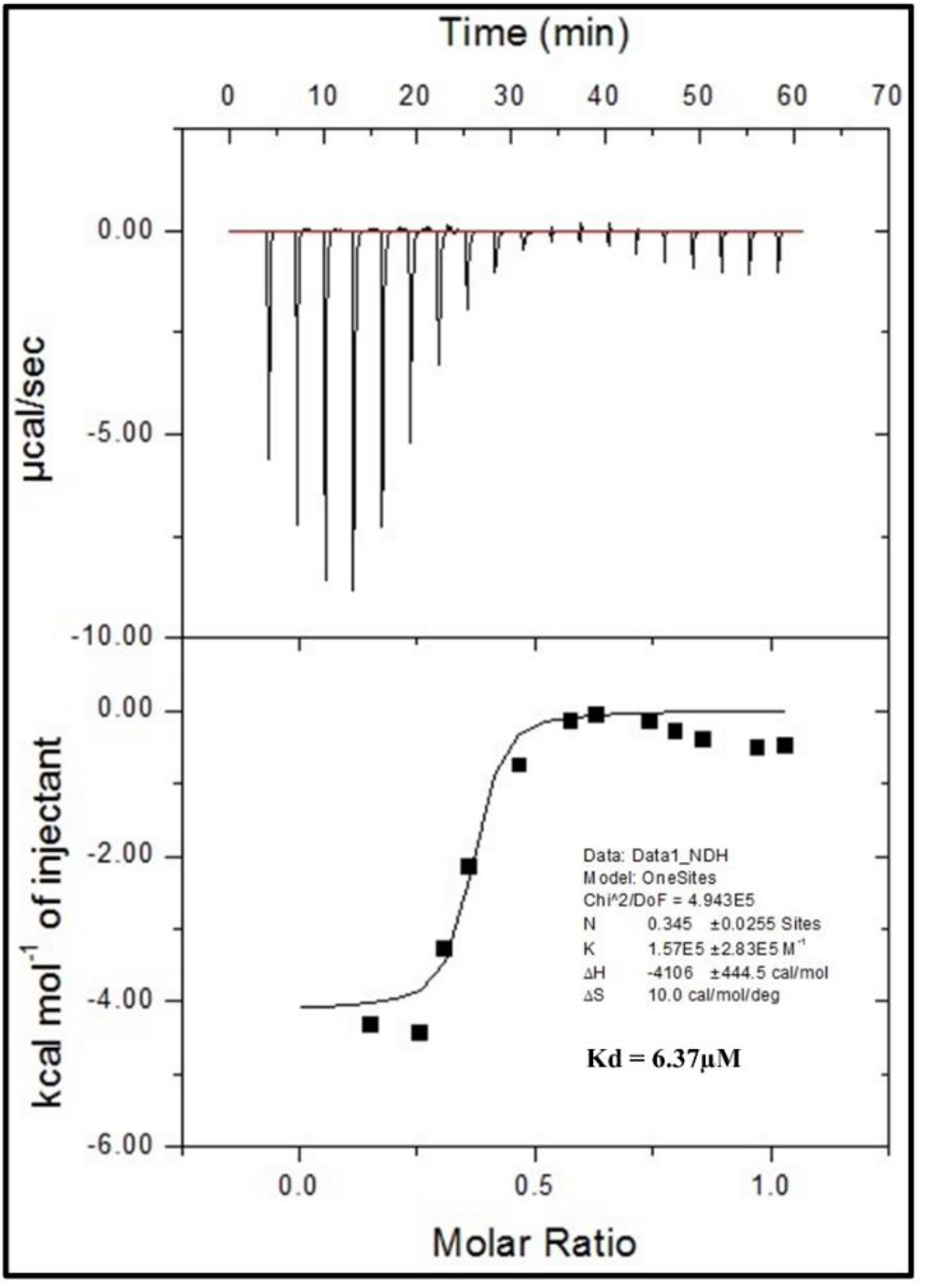
Isotherm binding of S100A1 to S100A4 at 25°C. The titration of S100A1 to S100A4 is shown in the top panel. The lower panel shows the incorporated data recovered after deducting the heat of dilution.

Fig 4 shows the binding iso-thermogram characterizing the S100A1-S100A4 interaction that describes the heat change that occurred when S100A1 was titrated with S100A4. S100A1-S100A4 binding was defined as free site model, yielding a stoichiometry of binding (1:1) We found the dissociation constant for the protein titration to be 6.37 μM.

### 3.5 Dissociation constant measurements based on fluorescence of the complex S100A1-S100A4

S100A1 protein consist one tryptophan residue at the sequence 90 and it is exposed. Tryptophan 90 is situated at the binding region of the S100A1. However, the excitation frequency of the tryptophan absorption band was found at 295 nm, and drop in fluorescence intensity shows the polarity change around W90. Decrease in intensity was observed after addition of S100A4 at the titration curve. These data were processed to get linear curve fitting and slope was calculated to get the Kd. Dissociation constant for S100A1-S100A4 complex was calculated which is in the range of about 10 μM. In Fig 5 we show the data graphed as 1/[S100A4] versus 1/(I - I_0_).

**Fig 5.**
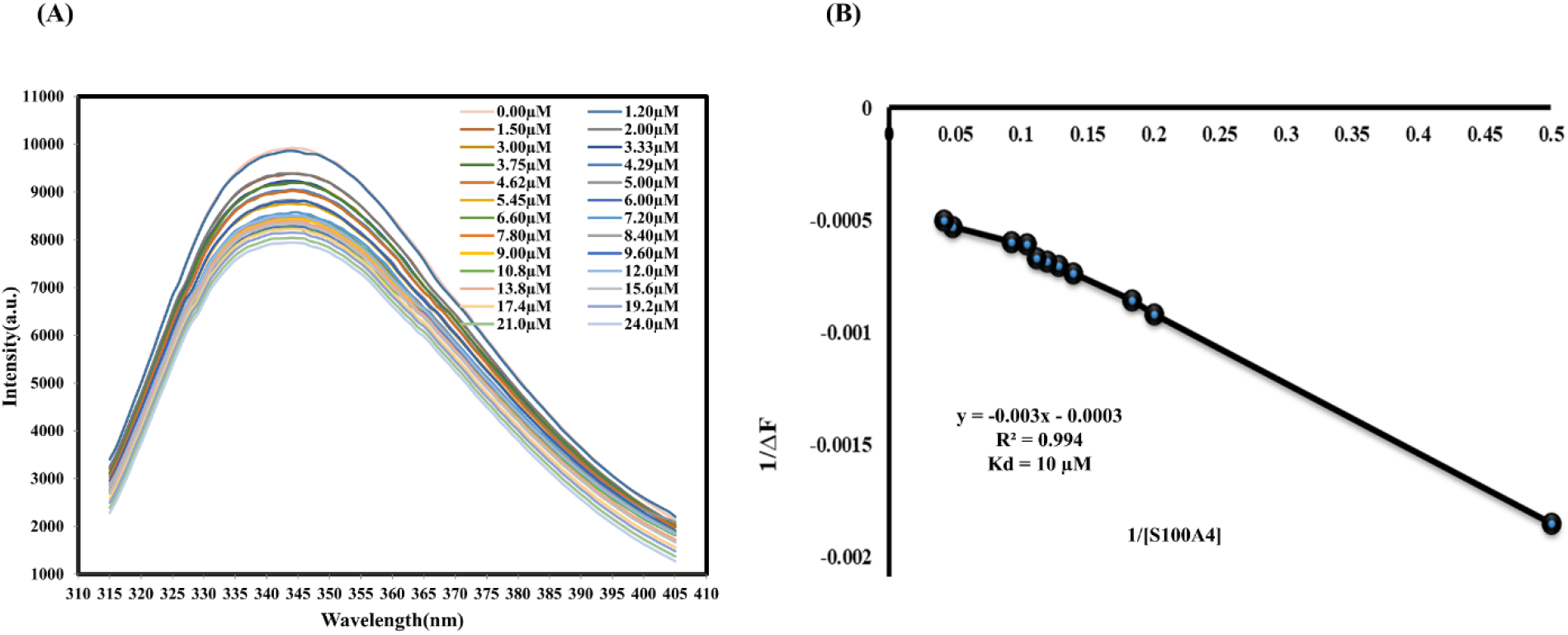
Emission of fluorescence spectra of S100A1 titrations showing a decrease in intensity with the addition of S100A4 at μM-level concentrations. (B) Change in fluorescence intensity versus S100A4 obtained at a wavelength of 351 nm. K_d_ was measured to be 10 μM using Equation 2 (62).

### 3.6 Functional bioactivity study with the WST-1 assay

A WST-1-based cell proliferation assay was used to describe the downstream signaling activity transmitted by the RAGE V domain through S100A1. We selected SW480 cells for the bioactivity functional assays because they have epithelial-like structure and are frequently used to study cancer *in vitro*. The SW480 cells were treated for 2 days with S100A1 prior to the variability analysis with the WST-1 assay.

Serum-starved SW480 cells were developed with S100A1 (concentration: 100 nM). We absorbed the 1.54-fold growth in the living cell count over without the serum control (Fig 6, Lane 2). This result can be clarified by the point that S100A1 interact with the RAGE V domain and hence induce cell proliferation (52). The proliferative capacity of the SW480 cells increased the number of cells by 1.57-fold when 100 nM S100A4 was added (Fig 6, Lane 3). A 1.38-fold decrease was observed in cells treated with both S100A1 (100 nM) and S100A4 (100 nM) proteins (Fig 6, Lane 4). These findings indicate that the S100A4 protein potentially inhibits contact between S100A1 and the RAGE V domain and inhibits cell proliferation activity.

**Fig 6.**
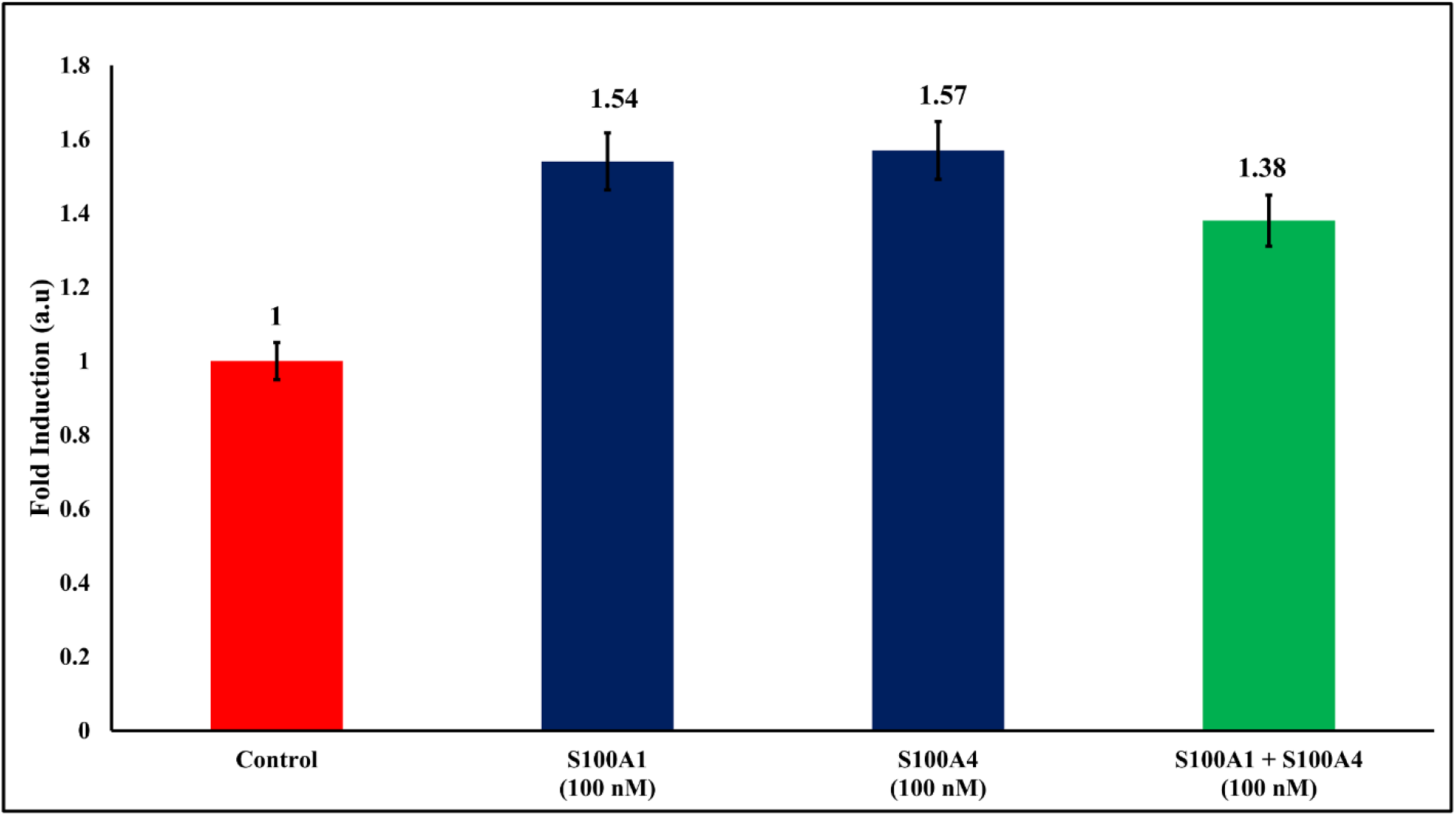
The SW480 cells were treated with 100 nM S100 A1 and S100A4 (blue, Lanes 2, 3) and 100 nM S100 A1 + S100A4 (green). The proportional cell counts after subsequent treatment with S100A1 are shown as fold inductions without serum media alone as the control (red).

The interactions between S100A1 and the V domain suggest that their binding may activate an auto-phosphorylation route that leads to several signal transduction cascades that regulate migration and cell proliferation. In this study, S100A1-RAGE V domain complex was used (Fig 7) (52).

**Fig 7.**
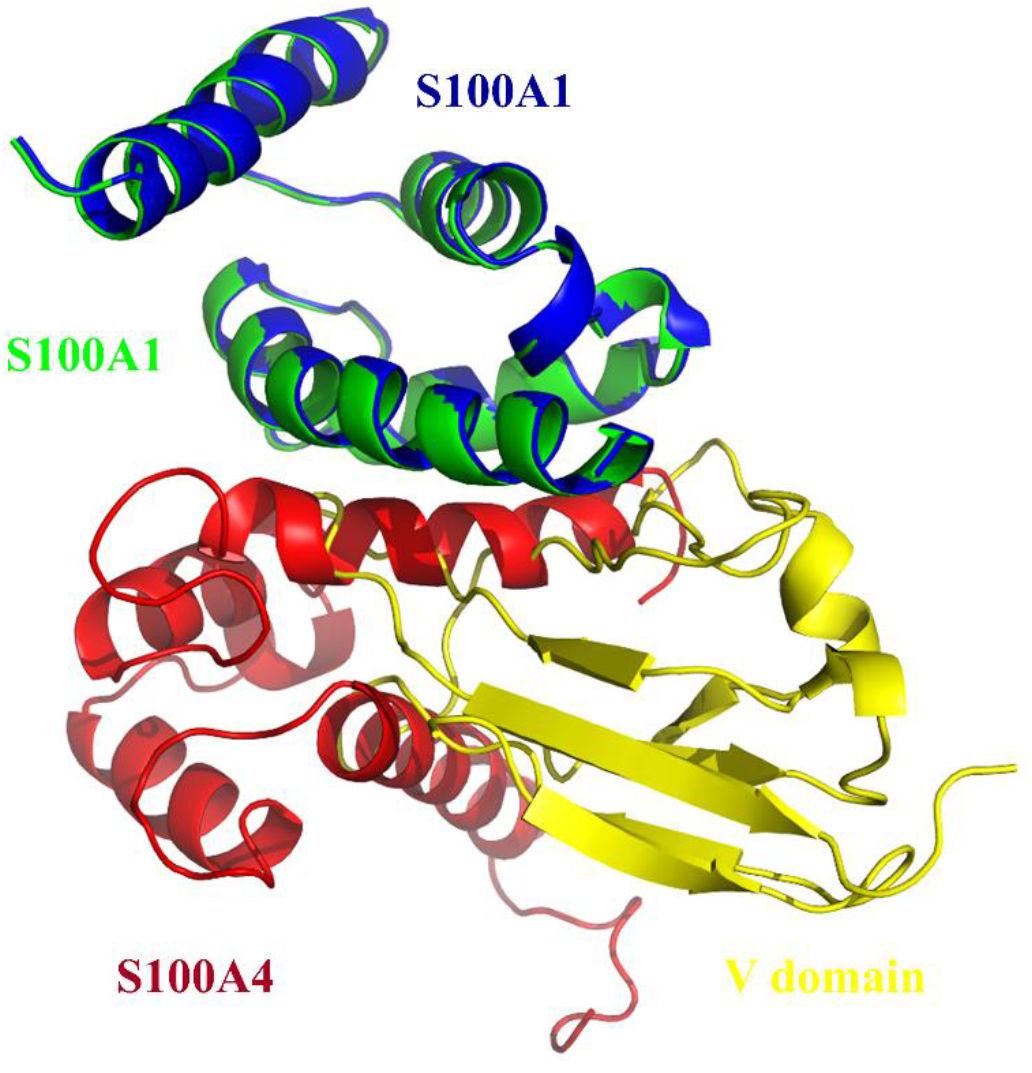
Overlapping structure of the S100A1-V domain complex (green, yellow) and the complex structure of S100A1-S100A4 (blue, red). S100A4 blocks interaction sites between S100A1-V domain complex.

## 4. Discussion

The paracrine, endocrine, and autocrine aspects of S100 family proteins are unknown. These proteins can be present in sufficient concentrations to determine cellular signaling in diverse cell types. We used NMR titration experiments to describe the binding between S100A1 and S100A4. Based on our results, we can conclude that S100A4 plays an essential role as an antagonist between the V domain and S100A1. Furthermore, our findings are essential to designing S100A4 analogs that may possibly act as effective blockers to inhibit the RAGE V domain-S100A1 route for the treatment of many kinds of human inflammation or cancer. The overlapping complexes shown below are (1) the S100A1 domain and (2) the S100A1-S100A4 complex (Fig 7). S100A1 and S100A4 are shown in blue and red, respectively, and the S100A1-V domains are shown in green and yellow.

The S100A4 molecule acts as an inhibitor to block the interaction of the V domain-S100A1 complex (52). This complex model may be valuable for improving antagonists and may potentially benefit protein-design studies that target S100A1 and the RAGE V domain.

## Acknowledgments

We acknowledge the Department of Chemistry, National Tsing Hua University, for allowing us to use their Varian NMR 700 MHz cryogenic spectrometer.

## Funding

This work was supported by the Ministry of Science and Technology (MOST) of Taiwan (grant number MOST 104–2113-M-007-019-MY3) and the China Medical University (grant number CMU104-S-02, Taiwan).

### Notes

The authors declare there are no conflicts of interest.

## Supporting Information

**Fig S1.**
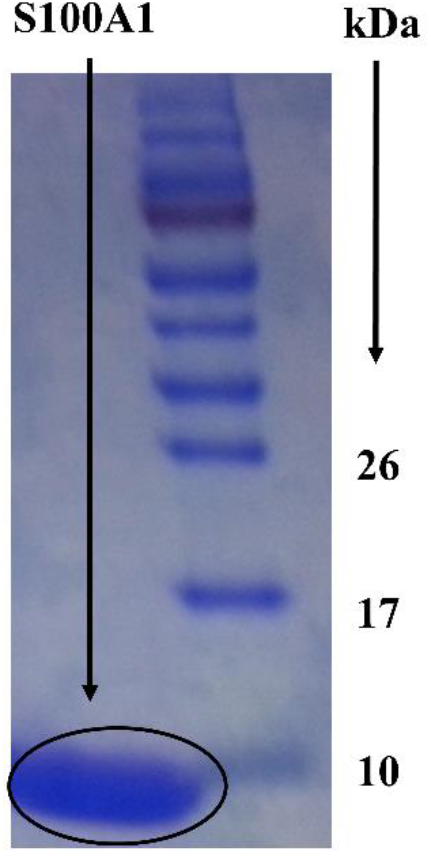
The SDS-PAGE band for the purified S100A1 protein showing a molecular weight of 10.5 kDa.

**Fig S2.**
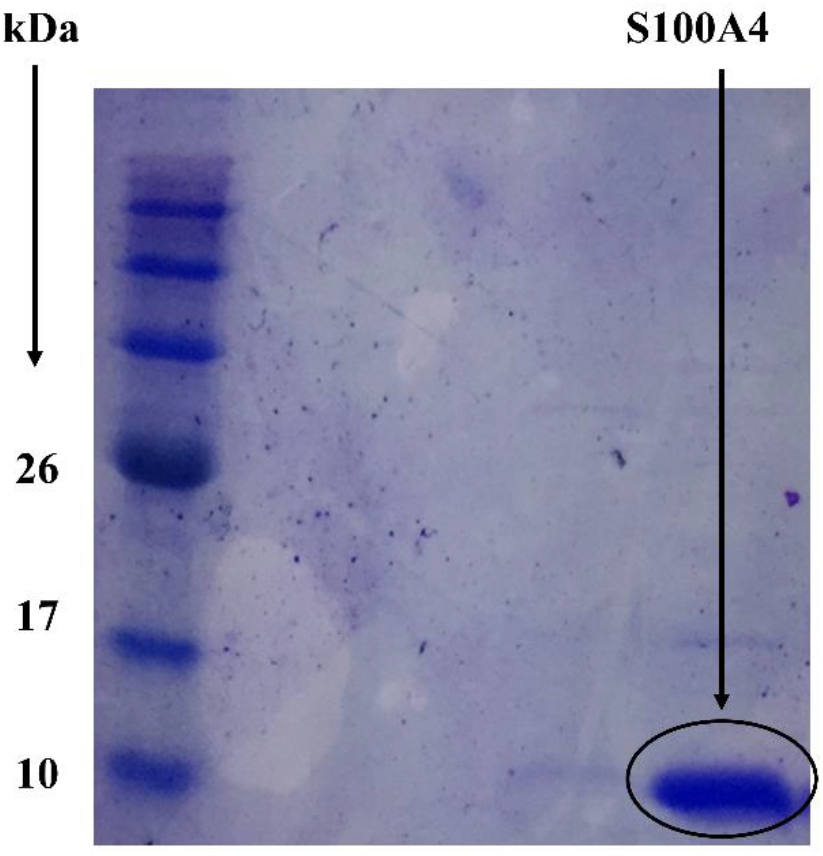
The SDS-PAGE band for purified S100A4 protein showing a molecular weight of 11.7 kDa.

**Fig S3.**
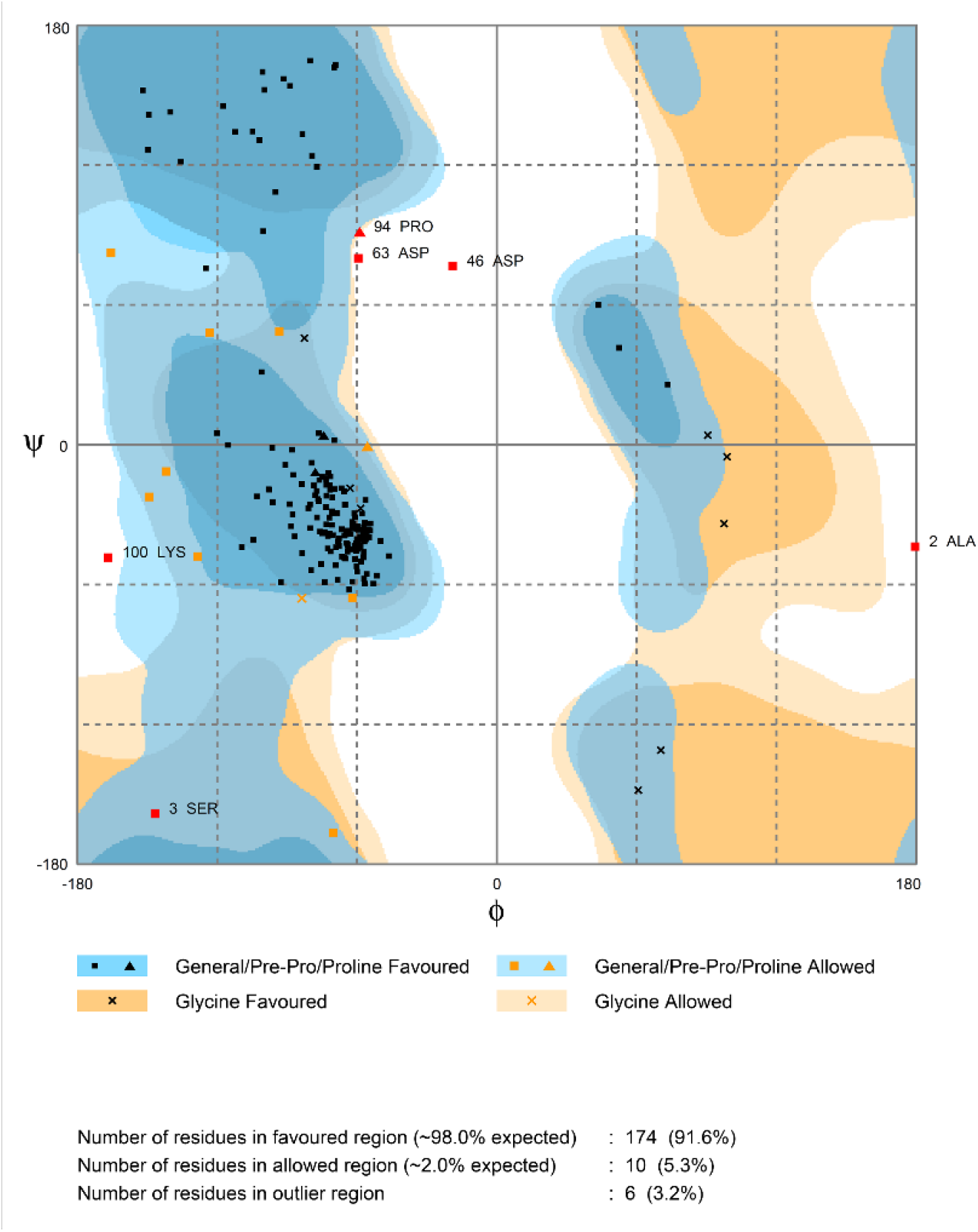
Ramachandran plot showing the complex of S100A1 with S100A4 according to the PROCHECK analysis. Ninety-one percent of the residues were in the favored area, 5.3% were in the allowed area, and 3.2% were in the disallowed region.

